# Brain-Wide Inferiority and Equivalence Tests in fMRI Group Analyses: Selected Applications

**DOI:** 10.1101/2021.04.22.440880

**Authors:** Martin Fungisai Gerchen, Peter Kirsch, Gordon Benedikt Feld

## Abstract

Null hypothesis significance testing is the major statistical procedure in the field of fMRI, but provides only a rather limited picture of the effects in a data set. When sample size and power is low relying only on strict significance testing may lead to a host of false negative findings. In contrast, with very large data sets virtually every voxel might become significant.

It is thus desirable to complement significance testing with procedures like inferiority and equivalence tests that allow to formally compare effect sizes within and between data sets and offer novel approaches to obtain insight into fMRI data. The major component of these tests are estimates of standardized effect sizes and their confidence intervals. Here we show how Hedge’s g, the bias corrected version of Cohen’s d, and its confidence interval can be obtained from SPM t maps. We then demonstrate how these values can be used to evaluate whether non-significant effects are really statistically smaller than significant effects to obtain “regions of undecidability” within a data set, and to test for the replicability and lateralization of effects.

This method allows the analysis of fMRI data beyond point estimates enabling researchers to take measurement uncertainty into account when interpreting their findings.

## Introduction

Functional magnetic resonance imaging (fMRI) relies heavily on statistical analyses to draw inferences and the use of null hypothesis significance testing (NHST) is the major statistical approach in the field. A major downside of the NHST framework is that it does not emphasize the comparison of effects, but rather pits a point null hypothesis against all alternatives, and thus provides only a rather restricted picture of the effects in a data set.

Without a priori constraints the large number of voxels that are recorded in fMRI inevitably leads to a multiple testing problem that is addressed by applying conservative corrections to the critical value and other methods for type I error control. However, this stringent correction can lead to studies that have low power (i.e., large type II error), especially when assuming weak distributed effects (Cremers et al., 2017). This is especially critical since small sample sizes tend to overestimate effect sizes in clusters passing the significance threshold, which leads to unreasonable expectations towards what constitutes a meaningful effect in fMRI research (Button et al., 2013; Ioannidis, 2008; Lindquist & Mejia, 2015; Reddan et al., 2017). This problem is especially critical when trying to replicate findings.

In contrast, with a very high number of participants an analysis will in most cases deliver a large number of significant voxels. In most cases it is also conceptually more interesting to know whether the activation of a brain region is more or less strongly associated with a specific behavior or intervention (see for example Bowring et al., 2019; Bowring et al., 2021).

To harness such information, it has been suggested for some time to supplement thresholded statistical parametric maps from NHST with maps of effect sizes (ES) (Jernigan et al., 2003). However, rather than relying just on point estimates for ESs, the construction of their confidence intervals (CIs) provides the means to conduct more formal equivalence testing (Lakens et al., 2018; see Figure 1 for an example explaining our approach using simulated data). Equivalence tests are able to show whether the data suggest that there is no effect larger (in absolute terms) than a specified threshold, the equivalence threshold. This can be achieved by using two one-sided tests, i.e., the first one-sided test is calculated against the positive equivalence threshold and the other one-sided test is calculated against the negative equivalence threshold. If both tests are significant, the measured effect is assumed to be between the two equivalence bounds. If only one of the bounds is used for testing, it is an inferiority test, as the procedure establishes that the effect is smaller than this bound. Importantly, instead of calculating one-sided tests against the equivalence bounds, a 90% confidence interval (CI) around the measured effect size can be calculated and equivalence is established if this confidence interval only contains values between the equivalence bounds.

**Fig. 1.**
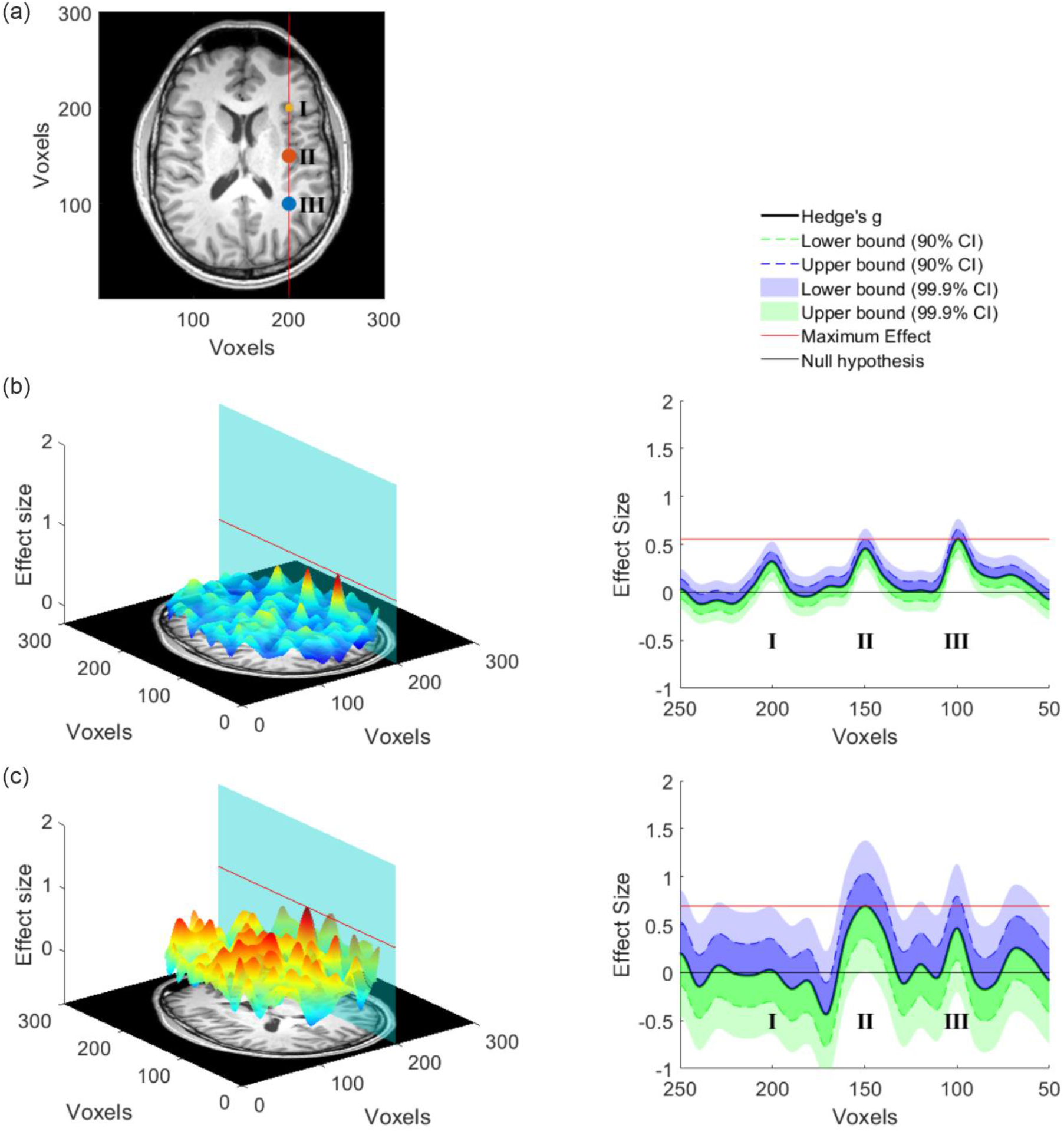
Data Simulation. (a) Three effects located at I, II and III (d = 0.28, d = 0.50 and d = 0.50, respectively) were generated for one fMRI-slice in a simulated dataset that compared two conditions (2 sample t-test, see supplement for details). (b) The left panel shows the effect size per voxel for a large sample (n = 500 per group) drawn from the simulated population. The plane cuts the 3-d graph at voxel 200, where the effects were inserted, and the red line marks the maximum effect size. In the right panel, all effect sizes lying on the plane are shown with the 90% and 99.9% CI added. The lower bound of the 99.9% CI in location I, II and III is above 0 indicating that these voxels would be significant in an uncorrected whole-brain t-test with α = 0.001. This means that in a large sample all three effects that were inserted into the data can be recovered. In addition, the maximum effect size (red line, effect at III) is larger than the 90% CI of the effect located I, which following the logic of equivalence testing, would enable to conclude that the effect in I is smaller than the effect in III. The same is not true for effects III and II as the red line cuts the 90% CI of effect II. (c) This panel shows the same as (b) however of a smaller sample (n = 50 per group). On the left it is evident that the effect sizes that are being estimated are much noisier, which is a result of the smaller sample size. On the right side it is evident that the CIs are also much enlarged, showing that the point estimate of the effect is much more uncertain. Consequently, only the effect in II is significant at the whole-brain threshold (p < 0.001). Importantly, we are also able to determine that most other voxels on this plane have effects that are smaller than the maximum effect found in the significant cluster of voxels. However, there is a cluster of voxels that are not significantly different from 0 at III, but that can also not be determined to be smaller than the effect present at II. Since the ground truth of the simulation is known this makes sense. Our method enables to identify such clusters in the whole brain and thereby allows deciding which brain areas can be excluded from being a relevant driver of certain behaviors and which cannot. Of course, the chosen threshold (peak voxel in this simulated case) will strongly influence the interpretation. Please see our use cases for indications on useful thresholds.

Such approaches have so far rarely been applied to MRI data (see Bowring et al., 2021; Pardoe et al., 2016; Reggev et al., 2020 for examples). With this paper we want to contribute to the use of this important statistical method in fMRI research. As a step towards this goal, we demonstrate here how Hedge’s g, the bias-corrected version of Cohen’s d, and its CI can be obtained using brain-wide t-maps from group analyses with one-sample and two-sample t-tests in statistical parametric mapping (SPM) with relative ease. We then provide concrete use cases that highlight how different inferiority and equivalence bounds allow different types of relevant inferences in fMRI data. We think that these procedures can provide a tool to further capture the richness of information in fMRI data and foster our understanding of brain processes.

## Methods

In this section we first discuss standardized ES and their CIs for t-tests in general and then specifically for fMRI group analyses.

### Standardized Effect Sizes for t-Tests

One of the best known ‘families’ of standardized ES is the ‘d’-family where a mean difference is standardized by a respective standard deviation

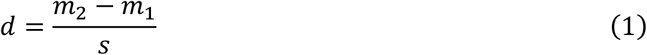

with subscripts referring to experimental groups. Different ‘flavors’ of d exist that differ in the exact estimation of the standard deviation. One specifically useful form for estimating effect sizes is based on the pooled standard deviation sp

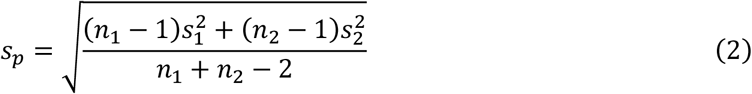

which gives d_p_

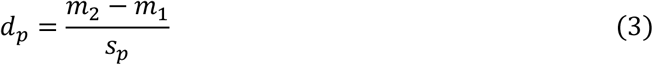

Please note that d_p_ is a version of Cohen’s d, but is also sometimes called Cohen’s g or Hedge’s g. Here we keep to the suggestion of Cohen (1988) to use subscripts for naming (see also Lakens, 2013). d_p_ is closely related to the t-test and can be directly calculated from t-values by

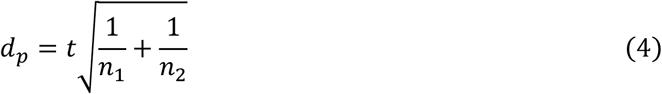

in the case of a two-sample t-test (Eq. (16.21) in Rosenthal (1994); Eq. (2) in Hentschke & Stüttgen (2011); Nakagawa & Cuthill, (2007), and equivalently by

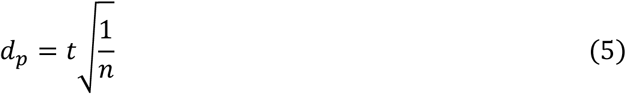

in the case of a one-sample t-test (see for example Bossier et al., 2019, p.16).

d_p_ is, however, a biased estimate of the population effect size, especially when it is based on a small sample size of n<20 per group, and needs to be corrected by a correction factor J (Hedges, 1981; Hedges & Olkin, 1985) to provide the unbiased effect size Hedge’s g

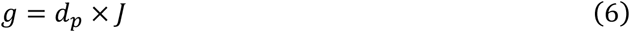

(Hedges, 1981) provides the following approximation to J by

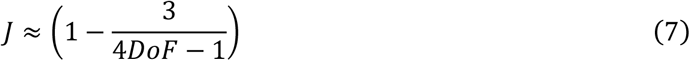

where DoF are the degrees of freedom used to estimate s.

CIs for ES can be estimated by bootstrap, exact analytical, or approximate analytical procedures (Hentschke & Stüttgen, 2011). Bootstrap and exact analytical procedures are computational much more demanding, while approximate analytical procedures are quite fast, but not always available (see Hentschke & Stüttgen, 2011). For the two-sample t-test an approximate analytical procedure is available which provides the standard error for the calculation of the CI of g

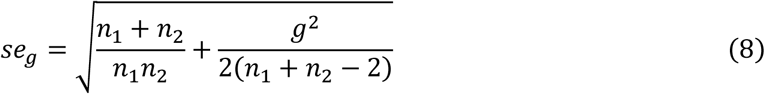

((Nakagawa & Cuthill, 2007), Eq. (17) in table 3).

In all cases, also when no approximate analytical procedure exists, CIs can be estimated by an exact analytical procedure based on the noncentral t-distribution (Smithson, 2003, p.34 ff.; Steiger & Fouladi, 1997). This procedure uses computationally intensive routines to estimate the CI for the noncentrality parameter of the noncentral t-distribution with noncentrality parameter Δ=t and the respective DoF. Because the cumulative distribution function of the noncentrality parameter is strictly increasing and monotonic, and the effect size is a monotonic, strictly increasing continuous function of this function, the obtained upper and lower limit of the noncentrality parameter CI can directly be inverted to the limits of the CI of the ES by

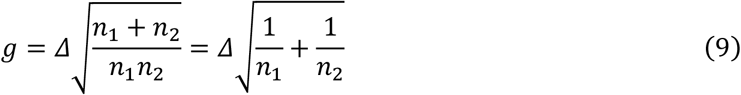

for the two-sample t-test ((Smithson, 2003), Eq. (4.7)) and by

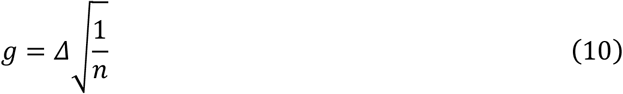

for the one-sample t-test (Smithson, 2003, Eq. (4.4)). Please see Steiger & Fouladi (1997) for a comprehensive explanation of the procedure. For estimating the limits of the CI of the noncentrality parameters we use the ‘ncpci.m’ function of the Measures of Effect Size Matlab toolbox (Version 1.6.1; https://github.com/hhentschke/measures-of-effect-size-toolbox) by Hentschke & Stüttgen (2011).

### Standardized Effect Sizes for t-Tests in fMRI

In fMRI analyses, however, t-tests are usually implemented in a general linear model (GLM) approach in which specific contrasts are tested for significance. Fortunately, the procedures based on “the noncentral t-distribution can be used to obtain confidence intervals for the standardized effect-size measure Cohen’s d in any situation where a t test is legitimate” (Smithson, 2003, p.62). However, unlike standard t-tests, in the GLM imaging analyses additional covariates, for example to correct for age and sex, are commonly included in the model and thus have to be taken into account when the procedures described above should be applied.

In SPM t-tests are implemented by

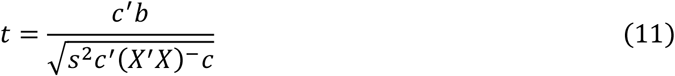

where X is the (pre-whitened and filtered) design matrix, c the contrast vector, b the estimated regression coefficients, c’ the transpose of c, (X’X)^-^ the pseudoinverse of X’X, and s^2^ the residual variance (see for example Penny et al., 2011, Eq. (8.12)). s^2^ is given by

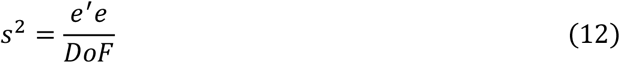

where e’e are the residual sum of squares and DoF=N-p where N is the number of samples (i.e. the number of rows of X) and p is the rank of X. These DoF are used in SPM to test for the significance of t.

The ES d for a specific contrast c in this case would have the form of

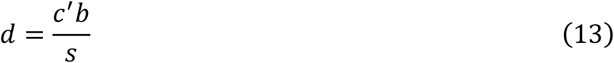

From Eq. (11) and Eq. (13), it follows that

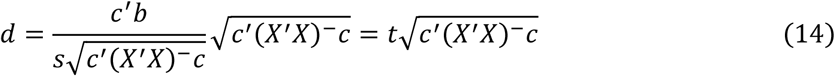

See also Bowring et al. (2021). Please note that for conventional one-sample t-tests 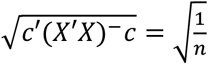 and DoF=n-1, and for conventional two-sample t-tests 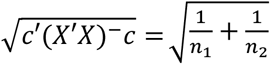 and DoF= n_1_-1+n_2_-1. In the case that covariates are added to the model, the DoF are decreased by the number of covariates and 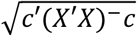 will take into account correlations of the covariates with the regressors included in the contrast.

Bias correction depends on the DoF and can be conducted in this case as in Eq. (6)

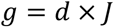

with the respective DoF entered in Eq. (7). Also, the estimation of the limits of the CI of the noncentrality parameter of the noncentral t-distribution depends on the DoF and can be conducted accordingly. The limits of the CI of the noncentrality parameter can then be converted to the limits of the CI of g by

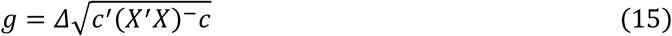

A script to estimate ES and their CI from SPM t maps is available on Github at https://github.com/Fungisai/g_ci_spm.

## Results

In this section we provide results for selected examples to demonstrate how the described methods can be used to obtain further insight into fMRI data.

### Within-Sample Comparison: “Maps of Undecidability”

Relying overly on NHST to separate activated from not activated brain regions in an (underpowered) fMRI study provides a distorted picture of the present effects. Here we suggest a formal strategy to address the question, which brain areas have ES that are statistically indistinguishable from the effects in a detected cluster above the statistical threshold. The resulting maps of such an analysis identify areas, which are “empty” in NHST, but where the suggestive conclusion of a smaller ES than in a detected cluster is not valid. Therefore, we call the obtained results “maps of undecidability”. More technically speaking, we test in every voxel whether the upper bound of its ES 90% CI is including or exceeding a reference ES representative for a detected cluster. Obviously, the results depend on the selected representative ES. In our example, we use the voxel with the median ES in the detected clusters as the reference.

We reanalyzed data from a monetary incentive delay task experiment performed by participants with Alcohol Use Disorder (AUD; nAUD=32) and healthy controls (HC; nHC=35) reported in Becker et al. (2017). We conducted analyses for the main effect (money > control) over both groups with a one-sample t-test (Figure 2) and for the group comparison (AUD > HC) with a two-sample t-test (Figure 3).

**Fig. 2.**
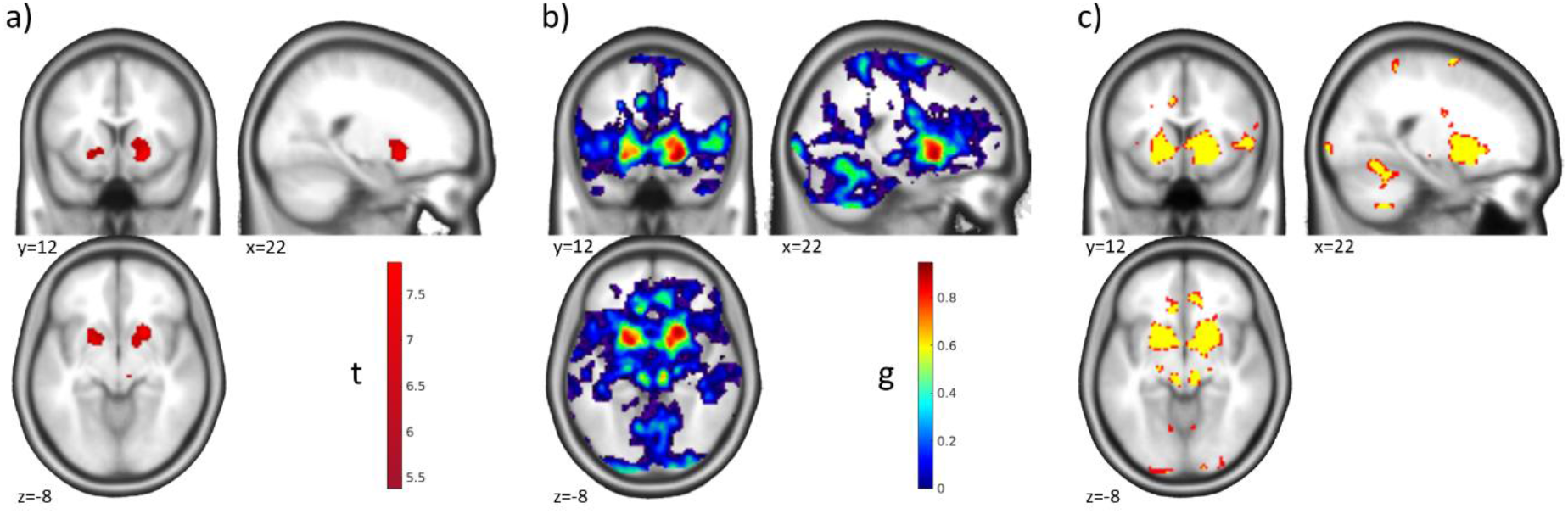
“Maps of Undecidability” – One-Sample t-Test. Results for a monetary incentive delay task in a sample of n=32 participants with Alcohol Use Disorder and n=35 healthy controls. a) Activation (p<0.05 whole-brain FWE corr.) for the main effect of the anticipation of monetary reward compared to the anticipation of verbal feedback in the whole sample showing a strong activation in bilateral striatum. b) Map of ES g for the activations shown in a). c) Areas of undecidability in yellow are marking voxels for which ES 90% CI included the median effect size (g=0.66) in the significant clusters. For comparison, uncorrected activation (p<0.001 unc.) is shown in red. In this specific example the areas of undecidability are largely overlapping with the uncorrected activation and are just slightly more spatially restricted. Please note that this correspondence depends on the exact chosen reference value and the properties of the specific data set for a given analysis. Reanalyzed data from Becker et al. (2017).

**Fig. 3.**
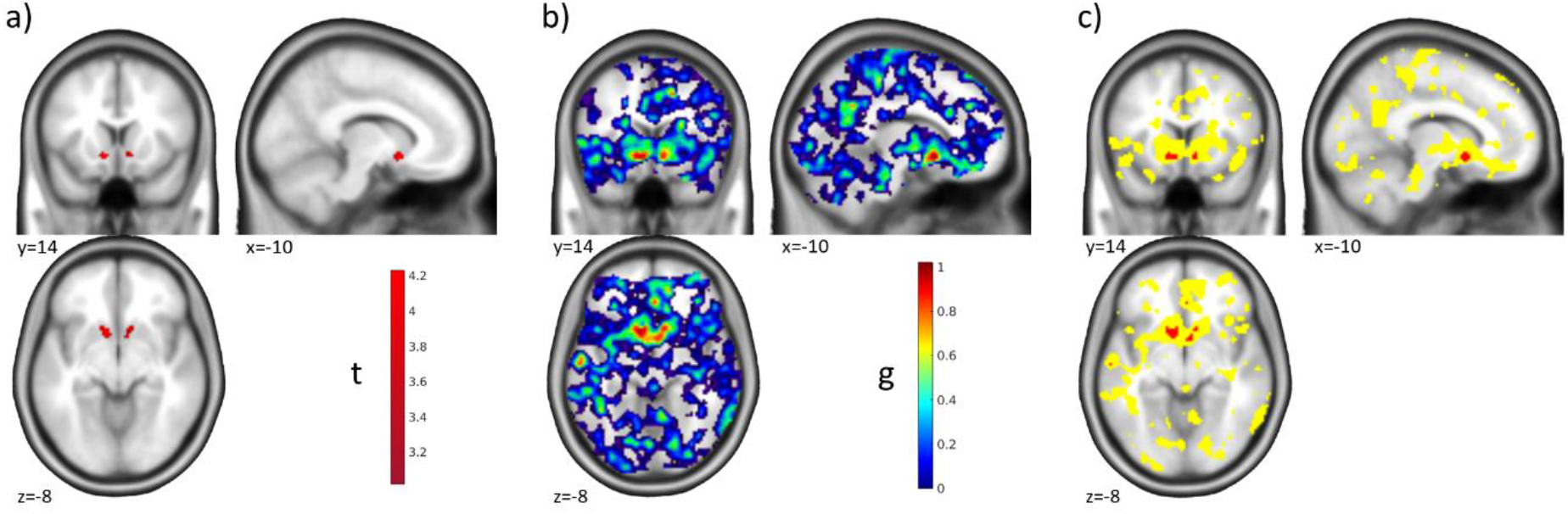
“Maps of Undecidability” – Two-Sample t-Test. Results of the group comparison for the monetary incentive delay task comparing participants with Alcohol Use Disorder and healthy controls. a) Activation for the group comparison (AUD>HC) based on ROI analyses in the left and right nucleus accumbens (p<0.025 FWE ROI analyses in each of the two ROIs). Participants with Alcohol Use Disorder showed stronger reactions in the nucleus accumbens than healthy controls. See Becker et al. (2017) for further details and discussion. b) Map of ES g for the activations shown in a). c) Areas of undecidability in yellow are marking voxels for which ES 90% CI included the median ES (g=0.7289) in the significant clusters. For comparison, uncorrected activation (p<0.001 unc.) is shown in red. In this example the uncorrected activation is very restricted and the areas of undecidability are rather large and extend well beyond. Reanalyzed data from Becker et al. (2017).

The one-sample t-test revealed a main effect of money > control in the bilateral ventral striatum at a threshold of p<0.05 whole-brain FWE corr. (Figure 2 a); corresponding ES in Figure 2 b)). “Undecidable” regions were for example found in the anterior cingulate cortex (ACC), right insula, and cerebellum (yellow in Figure 2 c)). Interestingly, in this example and at the chosen reference ES the undecidable areas were largely consistent with, and only minimally smaller than, the effect at p<0.001 unc. (red in Figure 2 c)).

In the group comparison the picture was quite different. Here, we identified a more localized effect by ROI analyses in the left and right nucleus accumbens at a threshold of p<0.025 FWE ROI analysis (p<0.05 corrected for two hemispheres; Figure 3 a); corresponding ES in Figure 3 b)). Here, this effect was largely consistent with the results at p<0.001 unc. (red in Figure 3 c)), and the areas of undecidability extended well beyond to, for example, striatum, ACC, posterior cingulate cortex, and insula (yellow in Figure 3 c)), reflecting a high uncertainty in the analysis about the uniqueness of the apparently very local effect. This result makes sense given the lower power of between-subjects comparisons to detect significant effects and corresponding larger confidence intervals of the effect sizes.

### Replication

Another directly apparent application of ES and their CIs in fMRI is testing for the replicability of detected effects. A voxel-wise strategy can be applied either with a general reference ES or with a reference map. For demonstration we reanalyzed a data set from an episodic memory task (Gerchen & Kirsch, 2017; see Supplement for further information) with two subsamples (N=136; n_1_=54, n_2_=82) scanned with the same protocol at different sites. Reflecting the situation that might occur in a replication study we use the smaller sample as the reference data set and the larger sample as the replication set. Both samples were originally analyzed together with the same analysis pipeline, which enables us to conduct voxel-wise comparisons. As replication criterion we tested whether the ES obtained with the reference sample fall into the voxel’s ES 90% CI in the replication sample. Following the usual approach in fMRI we focused on effects in one contrast direction (encoding > control), and restricted our analyses to voxels that had an ES g>0 in the reference data set. Similar tests could be added for the opposite contrast direction.

T maps thresholded at p<0.05 whole-brain FWE corrected for the two samples are shown in Figure 4 a) & b), the ES map for the reference sample is shown in Figure 4 c). The task leads to broadly distributed activations which are largely overlapping between the two samples. Interestingly, the voxel-wise test reveals further details beyond the overlap of significant effects (Figure 4 d)). First, small Es were replicated in large areas where no significant effect was detected. More importantly, very large ES in the original sample failed to replicate (red circles in Figure 4 d)), although the voxels were detected as significant in both samples, suggesting that the initial ES estimates in these areas were biased in the positive direction and should thus not be taken as representative of the underlying effect. This information would have been difficult to detect with NHST only.

**Fig 4.**
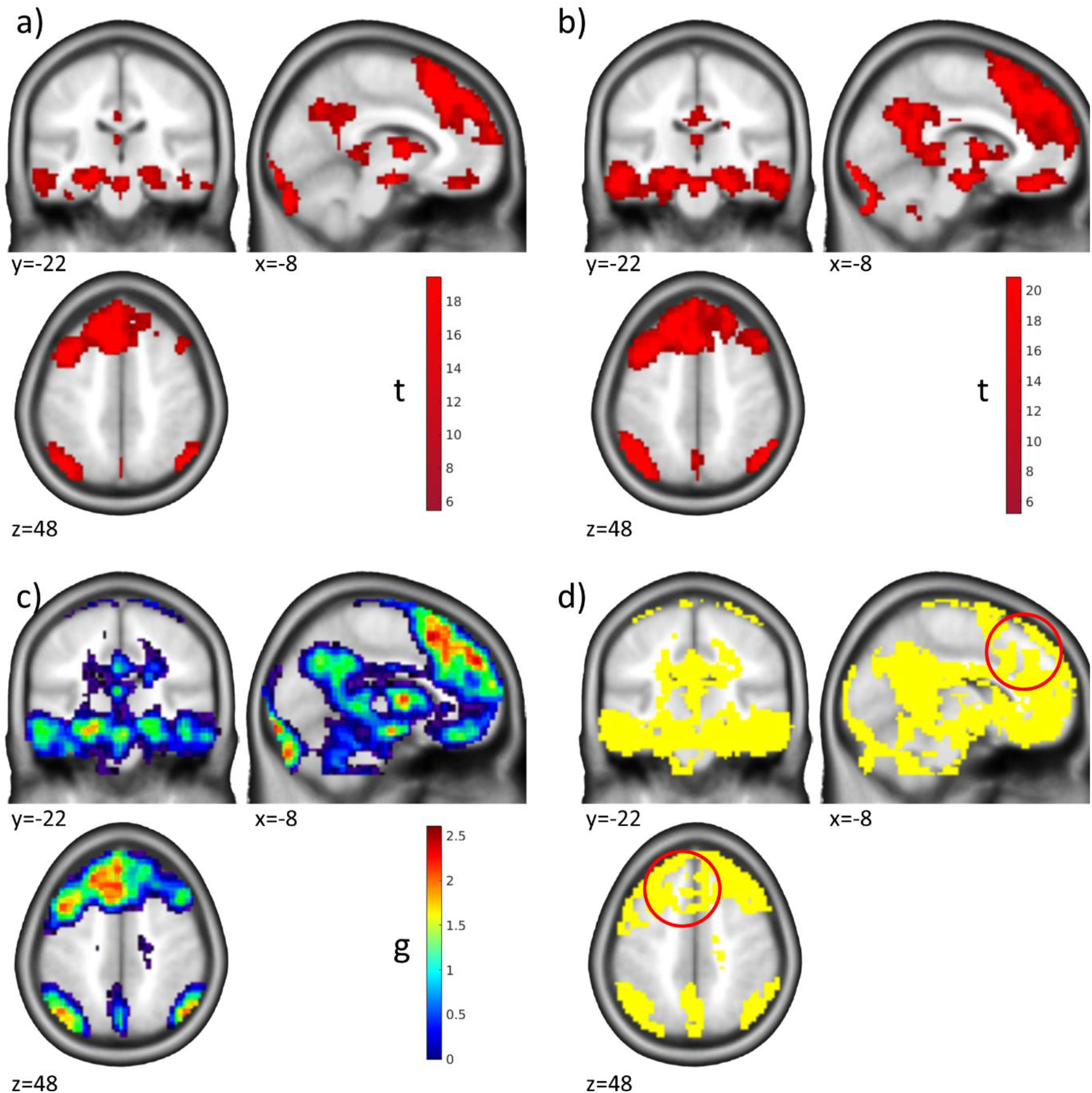
Replication of Effects. Results from the encoding phase of an episodic memory task are shown. a) Activation (p<0.05 whole-brain FWE corr.) for the contrast encoding > control in the reference sample of n1=54 participants. b) Activation (p<0.05 whole-brain FWE corr.) for the contrast encoding > control in the replication sample of n2=82 participants. Both samples were acquired in the same project with the same protocol but at different sites. c) Map of ES g for the activations shown in a). d) Yellow marks voxels where the ES 90% CI in the replication sample includes the ES of the voxel in the reference sample, which we define as a replication of the original effect size. Red circles: Area where the effect was significant in both samples but the reference ES did not replicate. Please note that only voxels are shown where the reference effect size was g > 0. Data from Gerchen & Kirsch (2017).

### Lateralization

An important question that arises in numerous contexts in functional neuroimaging is whether a detected effect is lateralized, i.e., more pronounced in one of the two hemispheres. Often a lateralization index is calculated (see for example Bradshaw, Bishop, et al., 2017; Bradshaw, Thompson, et al., 2017), but formal testing is difficult. Inferiority tests against a reference ES representative for a detected cluster offer a straightforward approach to address this question.

As an example, we use unpublished data from a statement judgement task where short written statements were presented to healthy right-handed participants (N=30) and rated as true or false (See Supplement for further information). Here, we did not focus on any specific experimental effect but analyzed the main effect of sentence presentation, which, beside others, showed strong activation in the left ventral occipito-temporal cortex and Broca’s area (Figure 5 a) & b)) associated with language processing (for example Bradshaw, Thompson, et al., 2017). Language processing has traditionally been described as lateralized to the left hemisphere in right-handed individuals (for example Bradshaw, Thompson, et al., 2017). Thus, we tested for these two clusters whether comparable effects are present in contralateral areas. For this we selected a reference ES reflecting a rather strong activation in the detected cluster and take the voxel with the 75th percentile ES in the reference cluster as the criterion. In other words, we test whether voxels whose ES 90% CI upper limit exceeds or includes the 75th percentile ES in the reference cluster are present in the respective contralateral region.

**Fig. 5.**
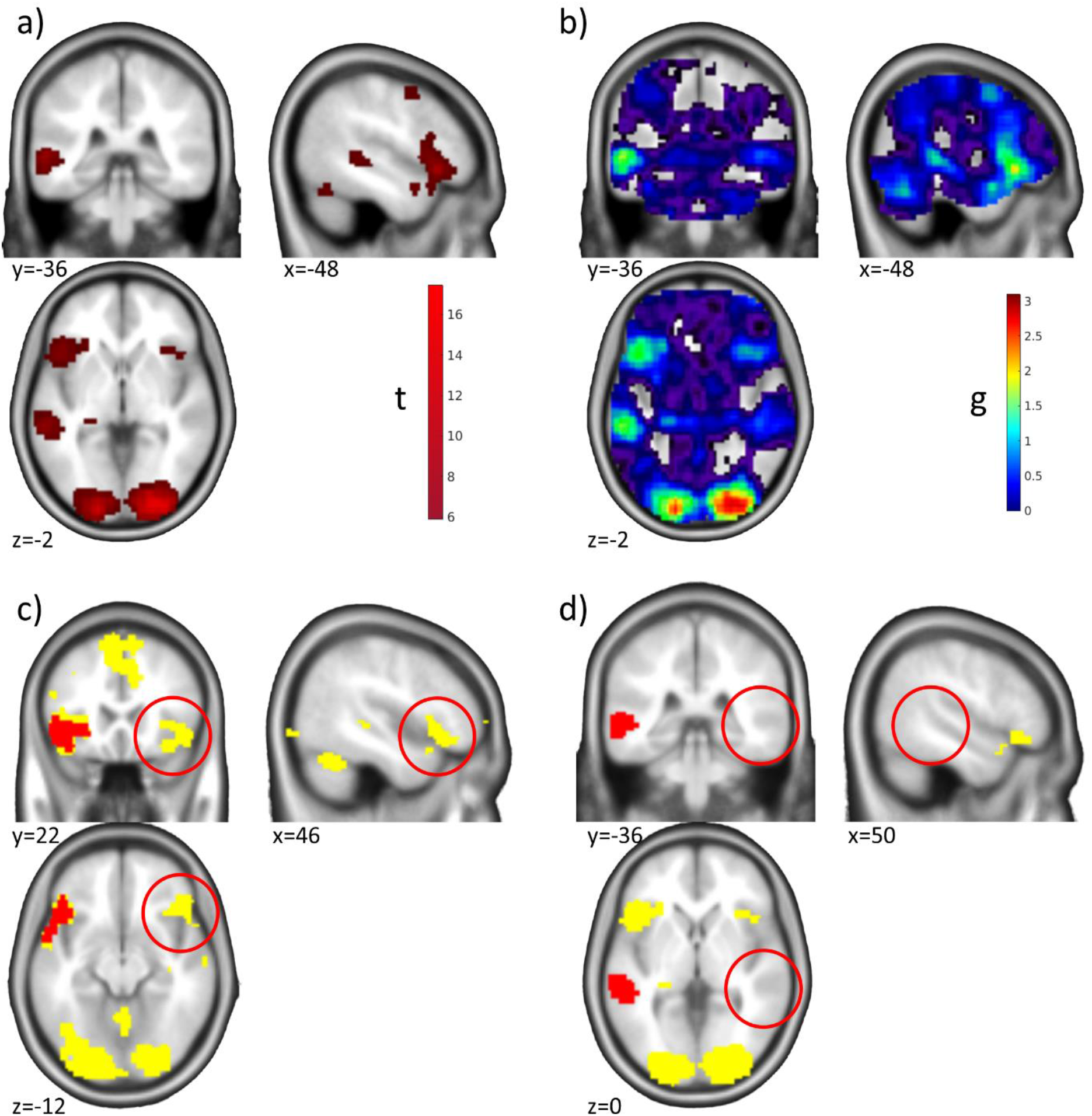
Lateralization of Effects. Results for a written statement presentation task are shown. a) Activation (p<0.05 whole-brain FWE corr.) for the main effect of written statement presentation in N=30 healthy participants. b) Map of ES g for the activations shown in a). c) Regions (yellow) that cannot be assumed to be smaller than the 75th percentile ES in the reference cluster (red) including Broca’s area. Red circles: Area in the right inferior frontal gyrus, suggesting contralateral effects in our data that cannot be shown to have a smaller effect than the reference cluster. d) Regions (yellow) that cannot be assumed to be smaller than the 75th percentile ES in the reference cluster (red) in the left ventral occipito-temporal cortex. Red circles: Inferior effects in the corresponding right left ventral occipito-temporal cortex, suggesting lateralization of effects in our data. Unpublished data by M.F. Gerchen.

For the FWE corrected significant reference clusters the 75th percentile ES was g=1.3242 for left Broca’s area, and g=1.454 in the left superior temporal cluster. Voxels with 90% ES CIs including the respective reference ES could be identified in the right inferior frontal cortex (Figure 5 c)), but not in the left ventral occipito-temporal cortex (Figure 5 d)). These results demonstrate how ES and their CIs can be used to provide evidence for, as well as against, lateralization in fMRI studies.

## Discussion

In this paper we described the construction of the standardized ES Hedge’s g and its CI for t-tests in statistical parametric mapping and demonstrated in selected examples how these can be used to identify “regions of undecidability”, to conduct voxel-wise replication tests, and for formal testing of lateralization of effects. Overall, our examples further demonstrate that NHST alone does not provide a conclusive picture about the effects contained in fMRI data, especially about the equivalence or inferiority of effects, and that complementary analyses as described allow important further insight into the results.

Obviously, a central decision for the described procedures with strong influence on the results and conclusions is the selection of the reference ES. Therefore, it is of uttermost importance that this selection is made a priori based on justifiable reasons related to the goal of the analysis and preregistered before the analysis is conducted. Within a data set, a number of possible criteria are for example the minimum, maximum, median, or quantile effect sizes in a reference cluster. As the ES are estimated in a voxel-wise manner, it might not be advisable to choose the mean or other summarizing values here.

It is further important to note that the smaller the reference ES (and the larger the CI) gets, the larger are the areas with overlapping CIs. It depends on the goal of the intended analysis what determines a liberal or conservative procedure. In our examples, we did not use a correction for multiple comparisons because the more conservative procedure was not to correct, and it remains an open question how a multiple comparison correction could be applied in the cases where it is deemed necessary.

If a data set should be used for replication of an external effect from another study, our procedures would for example allow to implement a voxel-wise small telescope approach in which one could tests for the existence of effects that an original study could have meaningfully examined (Simonsohn, 2015). Another interesting application for our method is using a smallest effect size of interest (SESOI) as the threshold (Lakens et al., 2018). In practice, it may be hard to determine the SESOI, but in large samples, it would allow excluding those voxels that have a significant activation that is too small to matter conceptually.

While our treatment covers only Hedge’s g as a specific ES for t-tests in SPM, these tests nonetheless cover a substantial part of analyses in the fMRI field. If other ES are needed, the MES toolbox (Hentschke & Stüttgen, 2011) provides a comprehensive library of ES and their CIs, which could be adapted for fMRI data in a similar way as we demonstrated here. Overall, we strongly believe the field of neuroimaging will benefit from providing evidence for absence of effects as much as for their presence and we hereby provide a method using a NHST-approach that can complement other approaches such as Bayesian statistics.

## Declaration of competing interest

The authors declare that they have no known competing financial interests or personal relationships that could have appeared to influence the work reported in this paper.

## Acknowledgements

The authors are grateful to Daniël Lakens, Peder Isager, and Thomas Nichols for helpful and insightful discussions.

## Funding information

This work was supported by the Deutsche Forschungsgemeinschaft (DFG, German Research Foundation) via an Emmy-Noether-Research-Group grant to GBF (FE 1617/2-1), and project funding to PK (SFB 636/ D6 and DFG Project-ID 402170461 – TRR 265 (Heinz et al., 2020). MFG was supported by the WIN programme of the Heidelberg Academy of Sciences and Humanities financed by the Ministry of Science, Research, and the Arts of the State of Baden-Württemberg. In addition, part of the work was funded by a grant from the German Ministry for Education and Research to PK (BMBF, 01GQ1003B). The funding sources had no involvement in study design, the collection, analysis and interpretation of data, and in the writing of the manuscript.

## Data and Code Availability

The fMRI data for the examples cannot be made publicly available due to protection of sensitive personal information. The Matlab script to estimate effect sizes and their CI from SPM t maps is available on Github at https://github.com/Fungisai/g_ci_spm.

## Supplemental Material

### Data simulation

We simulated one slice of fMRI data for a control group and an experimental group by setting up two 2d matrices with 0 for all voxels except for the three points indicated (I, II, III). For these points we chose the effect sizes of 0.45 (I), 0.8 (II) and 0.8 (III) for the experimental group’s matrix only. Next, we applied a spatial Gaussian filter, so that the effects at the three point were spread out spatially, which reduced the maxima to 0.28 (I), 0.50 (II) and 0.50 (III), i.e., the true underlying effects at those points. We then pulled random numbers for each simulated participant (N = 500 per group) using the numbers from the 2d matrices as mean per voxel and a standard deviation of 1 (therefore the simulated effects correspond to Cohen’s d). (b) The left panel shows the effect sizes (surface) for all voxels simulated within the underlying brain slice for a sample of n = 500 per group of the simulated data (Not: we did not constrain this simulation anatomically and the brain is purely for illustration purposes). We did not constrain our simulation anatomically, so no structural alignment should be expected. For Figure 1 we either analyzed the complete simulated data (b) or only a sub sample (c).

### Supplemental Materials & Methods

#### “Maps of Undecidability” Analysis

We reanalyzed data from Becker et al. (2017). The sample included nAUD= 32 participants with a diagnosis of Alcohol Use Disorder (Age 45.4±9 years; 77.4% male) recruited from the Central Institute of Mental Health inpatient addiction clinic and nHC=35 healthy control participants (Age 47.7±8.9 years; 65.7% male) matched for age, sex, and education recruited from the local population. Patients were abstinent for 11±5.6 days at the time of the experiment. All participants provided fully informed written consent and all procedures conformed to the Declaration of Helsinki and were approved by the local ethics committee of the Medical Faculty Mannheim of Heidelberg University.

In the experiment participants performed a monetary incentive delay reaction-time task (Kirsch et al. 2003). In this task participants have to react by pressing a button as fast as possible when a flash occurs. In each trial they are first informed by a cue about the consequences of a fast response (monetary reward: gain 2€; punishment avoidance: avoidance of a loss of 2€; verbal feedback: feedback on the performance; passive control condition: no response is required) which is shown until the flash occurs after 6s. After the response the participants see the respective feedback. The reaction time criterion is flexibly adapted to the reaction time of the participants. After the experiment participants received the money they had won in cash. Magnetic resonance imaging was conducted with a 3T Siemens Trio scanner (Siemens Healthineers, Erlangen, Germany) with a 12-channel head coil. T1-weighted anatomical images (MPRAGE) were collected with TR=2.3s, TE=3.03ms, flip angle 9° in 192 sagittal slices with slice thickness 1.0mm, in-plane resolution of 1mmx1mm and field of view FoV=256×256mm. 267 functional images were acquired with an echo-planar imaging (EPI) sequence with TR=2s, TE=30 ms, flip angle 80° in 28 slices with slice thickness 4.0 mm with 1mm gap and matrix size = 64×64.

Analyses were conducted with SPM8 (v4515) in MATLAB R2011b. The functional images were slice-time corrected, realigned, co-registered to the anatomical image, normalized into MNI space and smoothed with a 6mm full-width at half-maximum Gaussian kernel. The ART toolbox (http://www.nitrc.org/projects/artifact_detect) was used to identify motion-affected volumes with volume-to-volume movements >1mm and global intensity changes z>7.

First-level analyses were conducted with general linear models including the event-related condition regressors convolved with the canonical HRF, 6 conventional motion parameters, white matter (WM) and cerebrospinal fluid (CSF) signals and the ART dummy regressors as covariates. A high-pass filter with a cut-off period of 128s was applied. Only the contrast monetary reward > verbal feedback was used and the contrast maps of the participants were used for the second-level analysis. A one sample t-test was conducted for the main effect of monetary reward > verbal feedback at a significance threshold of p<0.05 whole-brain FWE corr. ROI analyses in the left and right nucleus accumbens at a threshold of p<0.025 FWE ROI corr. (p<0.05 corrected for two hemispheres) was conducted with a two sample t-test to test for group differences in the direction AUD>HC. In contrast to the analyses reported in Becker et al. (2017) we used here a slightly more robust procedure to conduct the ROI analyses but come to the same results.

#### Replication Analysis

We reanalyzed data from Gerchen & Kirsch (2017). The data set comprised two samples with n_1_=54 and n_2_=82 healthy right-handed participants. The average age in the whole sample was 31.99±9.72 years and 47.8% of the participants were female.

In the experiment participants performed an episodic memory task (Erk et al., 2010) in which they learned associations between faces and professions. Here, we analyzed only data from the encoding phase. In this phase participants saw 16 face-profession pairs (each twice) in an ABAB block design with 4 experimental and 6 control blocks and were instructed to remember the associations. During face presentation participants had to indicate by a left or right button press whether faces and professions were matching well in their own opinion. During the control blocks schematic heads were shown and the participants indicated which of the two ears was larger.

All participants provided fully informed written consent and all procedures conformed to the Declaration of Helsinki and were approved by the local ethics committees of the Medical Faculties Mannheim and Heidelberg of Heidelberg University.

Magnetic resonance imaging was conducted with two similar 3T Siemens Trio scanners (Siemens Healthineers, Erlangen, Germany) at the Central Institute of Mental Health Mannheim and at the University of Heidelberg, Germany. 244 functional images were acquired with echo-planar imaging (EPI) with TR=1.8s, TE=30ms, flip angle 73°, in 33 slices with slice thickness 3.0 mm with 1mm gap, a field of view of FoV=192 mm, and GRAPPA with iPAT=2. Analyses were conducted with SPM8 (v4667) in MATLAB R2011b. The functional images were slice-time corrected, realigned, normalized into MNI space, rescaled to 3×3×3mm, and smoothed with a 6mm full-width at half-maximum Gaussian kernel. The ART toolbox was used to identify motion-affected volumes with volume-to-volume movements >0.5mm and global intensity changes z>6.

First-level analyses were conducted with general linear models including the block regressors convolved with the canonical hemodynamic response function (HRF), 6 conventional motion parameters, the CSF signal and the ART dummy regressors as covariates. A high-pass filter with a cut-off period of 128s was applied. Only the contrast encoding > recall was used and the contrast maps of the participants were used for the second-level analysis which were conducted with one sample t-tests within each subsample with age and sex as covariates. A significance threshold of p<0.05 whole-brain FWE corr. was applied.

#### Lateralization Analysis

For the lateralization analysis we used unpublished data from a written sentence presentation and judgement task by M.F. Gerchen. The sample consisted of N=30 healthy right-handed participants (age 23.37±1.43 years; 73.3% female) who were mainly university students. In the task short written statements with a maximum of 10 words from the categories religion, facts, conspiracy theories, superstition, and politics were presented for a duration of 6.56-8.2s in 100 trials. After each statement participants answered the forced-choice question whether they think the statement is true or not with a button press and rated their certainty on a visual analog scale from 0-100%. All participants provided fully informed written consent and all procedures conformed to the Declaration of Helsinki and were approved by the local ethics committee of the Medical Faculty Mannheim of Heidelberg University (2019-724N).

Magnetic resonance imaging was conducted with a 3T Siemens Trio scanner (Siemens Healthineers, Erlangen, Germany). T1-weighted anatomical images (MPRAGE) were collected with TR=2.3s, TE=3.03ms, flip angle 9° in 192 sagittal slices with slice thickness 1.0mm, in-plane resolution of 1mmx1mm and field of view FoV=256×256mm. Functional images were acquired with an echo-planar imaging (EPI) sequence with TR=1.64s, TE=30ms, flip angle 80° in 30 slices with slice thickness 3.0 mm with 1mm gap with 3×3mm in-plane resolution, and GRAPPA with iPAT=2. The duration of the experiment depended on the response speed and the average scan duration was 866.1±65.82 volumes. During scanning pulse and respiration were monitored and saved.

Analyses were conducted with SPM12 (v7219) in MATLAB R2017a. The anatomical image was segmented and normalized to MNI space. The functional images were slice-time corrected, realigned, co-registered to the anatomical image, normalized into MNI space, rescaled to 3×3×3mm resolution and smoothed with an 8mm full-width at half-maximum Gaussian kernel. The ART toolbox was used to identify motion-affected volumes with volume-to-volume movements >0.5mm and global intensity changes z>4. The TAPA PhysIO toolbox (Kasper et al., 2017) was used to estimate physiological nuisance regressors.

First-level analyses were conducted with general linear models including the event-related condition regressors convolved with the canonical HRF, 6 conventional motion parameters, WM, CSF, and whole-brain signals were used as covariates along with the ART dummy regressors and the PhysIO physiological nuisance regressors. A high-pass filter with a cut-off period of 128s was applied. Here we only analyzed the main effect of written sentence presentation. The reported analyses were not intended when the study was planned and are not part of the main results of the study. The contrast maps of the participants were used for second-level analysis with a one sample t-test in which age and sex were included as covariates. A significance threshold of p<0.05 whole-brain FWE corr. was applied.

## Notes

https://github.com/Fungisai/g_ci_spm

### Competing Interest Statement

The authors have declared no competing interest.

